# HSP90 inhibitors disrupt a transient HSP90-HSF1 interaction and identify a noncanonical model of HSP90-mediated HSF1 regulation

**DOI:** 10.1101/207001

**Authors:** Toshiki Kijima, Thomas L. Prince, Megan L. Tigue, Kendrick H. Yim, Harvey Schwartz, Kristin Beebe, Sunmin Lee, Marek A. Budzynski, Heinric Williams, Jane B. Trepel, Lea Sistonen, Stuart Calderwood, Len Neckers

## Abstract

Heat shock factor 1 (HSF1) initiates a broad transcriptional response to proteotoxic stress while also mediating a cancer-specific transcriptional program. HSF1 is thought to be regulated by molecular chaperones, including Heat Shock Protein 90 (HSP90). HSP90 is proposed to sequester HSF1 in unstressed cells, but visualization of this interaction *in vivo* requires protein crosslinking. In this report, we show that HSP90 binding to HSF1 depends on HSP90 conformation and is only readily visualized for the ATP-dependent, N-domain dimerized chaperone, a conformation only rarely sampled by mammalian HSP90. We have used this mutationally fixed conformation to map HSP90 binding sites on HSF1. Further, we show that ATP-competitive, N-domain targeted HSP90 inhibitors disrupt this interaction, resulting in the increased duration of HSF1 occupancy of the *hsp70* promoter and significant prolongation of both the constitutive and heat-induced HSF1 transcriptional activity. While our data do not support a role for HSP90 in sequestering HSF1 monomers to suppress HSF1 transcriptional activity, our findings do identify a noncanonical role for HSP90 in providing dynamic modulation of HSF1 activity by participating in removal of HSF1 trimers from heat shock elements in DNA, thus terminating the heat shock response.

## Introduction

Heat shock factor 1 (HSF1) is an evolutionarily conserved transcription factor that initiates the cytoprotective heat shock response (HSR). Found throughout the eukaryotic kingdom, HSF1 allows for the cellular adaptation to proteotoxic stress^1^. Through an incompletely defined mechanism, mammalian HSF1 monomers in cytosol are activated and form trimers, translocate into the nucleus, and bind sequences of DNA known as heat shock elements (HSE), ideally represented as nGAAnnTTCnnGAAn^2 3^. Throughout this process HSF1 is heavily post-translationally modified and interacts with numerous cellular components. The binding of HSF1 trimers to HSE induces the transcription of a specialized set of genes known as molecular chaperones while also repressing the expression of other genes^4 5^, although the repressive effect of HSF1 is controversial^6^. Once expressed, these molecular chaperones (or Heat Shock Proteins, HSPs) act to stabilize the three-dimensional structure of numerous cellular proteins, thus helping to maintain cellular proteostasis. HSP90 and HSP70 are ATP-dependent HSPs that interact with a large sector of the eukaryotic proteome while also modulating HSF1 transcriptional activity^7 8 9^. The relationship between HSF1 and one or more components of the cellular proteostasis network is thought to represent a primary axis in the control of the HSR ^7 10^.

In cancer, HSF1 enables malignant cell growth and is overexpressed in a number of tumor types associated with poor prognosis^11 12 13^. Although HSF1 does not initiate oncogenic transformation, tumors become addicted to HSF1 activity as their microenvironments become increasingly toxic and as they require higher levels of HSPs to maintain proteostasis^14^. Moreover, many oncogenes that drive tumorigenesis are metastable and rely on HSPs to sustain their activity. This is particularly true for mutated or overexpressed kinases and transcription factors that interact with HSP90^15^. HSF1 also promotes a cancer-specific transcriptional program that supports malignancy through the expression of genes for proliferation, anabolic metabolism, metastasis and apoptosis prevention. Comprised of over 500 genes, this cancer-specific HSF1 transcriptome is associated with poor clinical outcomes ^11^.

The human *HSF1* gene is encoded on chromosome 8q24 by 14 exons that produce two splice variants. The largest variant, which is described in this report, is translated into 529 amino acids. HSF1 has a predicted molecular weight of 57 kDa, yet migrates at approximately 75 kDa on SDS-PAGE due to a large number of post-translational modifications (PTMs), including phosphorylation, acetylation and sumoylation^16 17^. The overall structure of HSF1 is mostly disordered except for the evolutionarily conserved N-terminal DNA-binding domain (DBD) that forms a winged helix-turn-helix structure^18 19^. The rest of HSF1 is predicted not to maintain a stable tertiary structure, a feature observed for many proteins involved in transcription and cellular regulation^20^. Following the DBD and a linker region is the set of heptad repeats (HR-A/B) that form the leucine zippers that allow for HSF1 trimerization^21^. Adjacent to the HR-A/B, the unstructured regulatory domain (RD) is the molecular region understood to be capable of sensing heat and initiating the HSR^22^. The RD contains numerous phosphorylation sites^23^ and functions, along with a portion of HR-A/B, to repress the transcriptional activity of NF-IL6^24^. Another heptad repeat (HR-C), C-terminal to the RD, is understood to sequester HR-A/B in an intramolecular interaction that suppresses spontaneous HSF1 trimerization^21^. Most recently, Hentze et al have shown that HSP90β interacts with the HR-C domain in solution to promote its temperature-sensitive dissociation from HR-A/B, thus reducing the temperature at which HSF1 is able to trimerize^25^. The C-terminal transactivation domain (TAD) is highly disordered and is required for robust HSF1 transcriptional activity in response to stress ^26^. The TAD is also a site of HSP70 binding, which negatively regulates HSF1 transcriptional activity^27^.

While the interaction of HSP70 with HSF1 in unstressed cells is readily observed by co-immunoprecipitation^27^, interaction of endogenous HSP90 and HSF1 is not readily detectable and requires cross-linking to stabilize the association^7^. These data argue against a frequently cited model postulating sequestration of HSF1 monomers by HSP90 under non-stress conditions (7), but they would support such a role for HSP70, as has recently been proposed in yeast^28^. Chaperone binding to HSF1 monomers is thought to prevent spontaneous trimerization. In such a model, HSF1 is released from chaperone suppression and allowed to trimerize when cells experience severe proteotoxic stress that generates a large population of denatured client proteins and an increased demand for chaperones^29^. Other studies have shown that HSP90 can also bind the trimerized DNA-bound state of HSF1 and that this interaction attenuates HSF1 transcriptional activity^8^. Thus, HSP90 has variously been proposed to function as a sensor, repressor and/or attenuator of the HSR.

Determining the mechanism by which HSP90 affects the HSR is of medical significance. Currently, HSP90 is the only HSP whose targeting by small molecules has been extensively evaluated in the clinic as a strategy to treat cancer^30^. However, although HSP90 inhibitors have shown great promise in pre-clinical studies due to their ability to disrupt proteostasis and to promote the degradation of oncogenic kinases and other pro-growth components^31^, their single agent activity in patients has not been impressive^32 33^. This discrepancy has been ascribed, in part, to the observation that ATP-competitive HSP90 inhibitors promote the HSR and result in the increased expression of HSP70, HSP27, and other pro-survival components^34 35^, possibly offsetting any anti-cancer benefit of these drugs. Moreover, HSP90 inhibitors may enhance the recently identified cancer-specific HSF1 transcriptional program^11^. How this is mechanistically achieved in the absence of robust HSP90/HSF1 interaction has remained obscure.

In this report, we define the interactions of HSF1 with both nuclear-cytosolic paralogs of HSP90, stress-inducible HSP90α and constitutively expressed HSP90β, and with endogenous HSP70.Remarkably, we readily observed interaction of HSF1 with mutant HSP90 trapped in an ATP-dependent “closed” conformation (unable to hydrolyze ATP), and we found this interaction to be disrupted by N-terminal HSP90 inhibitors that are currently being clinically evaluated. To determine the domains in HSF1 that bind HSP90 and HSP70 we made a series of HSF1 deletion mutants and analyzed their interactions by immunoprecipitation-western blot. We observed that the binding of HSP90α, HSP90β and HSP70 uniquely map across HSF1. This was complemented by analysis of the effects of these deletion constructs on HSF1 trimerization and HSR reporter activity. Finally, we found that HSP90 inhibitors disrupt association of “closed” HSP90 proteins with HSF1, while concomitantly increasing the duration of HSF1 binding to *hsp70* promoter and extending the duration of heat-induced HSF1 transcriptional activity. While our data do not support a role for HSP90 in sequestering HSF1 monomers, our findings reveal that HSP90 inhibitors interfere with a noncanonical role for HSP90 in providing dynamic modulation of HSF1 activity by removing HSF1 trimers from heat shock elements in DNA.

## Results

### Wild type HSF1 readily interacts with N-domain dimerized (“closed” conformation) HSP90

Previous studies have suggested that the intracellular interaction of HSF1 and HSP90 is weak and transient, and requires chemical crosslinking for visualization^7^. We showed recently that the strength of this interaction may be influenced by HSP90 conformation^36^. Binding of ATP to N-terminal domains of HSP90 promotes their transient dimerization and induces a “closed” conformation that is poised for ATP hydrolysis. ATP binding and hydrolysis, in turn, drive the structural rearrangements in HSP90 that support chaperone function. Upon release of ADP, or in the presence of ATP-competitive HSP90 inhibitors, HSP90 N-domains revert to an undimerized state (“open” conformation) (Figure 1A).

**Figure 1:**
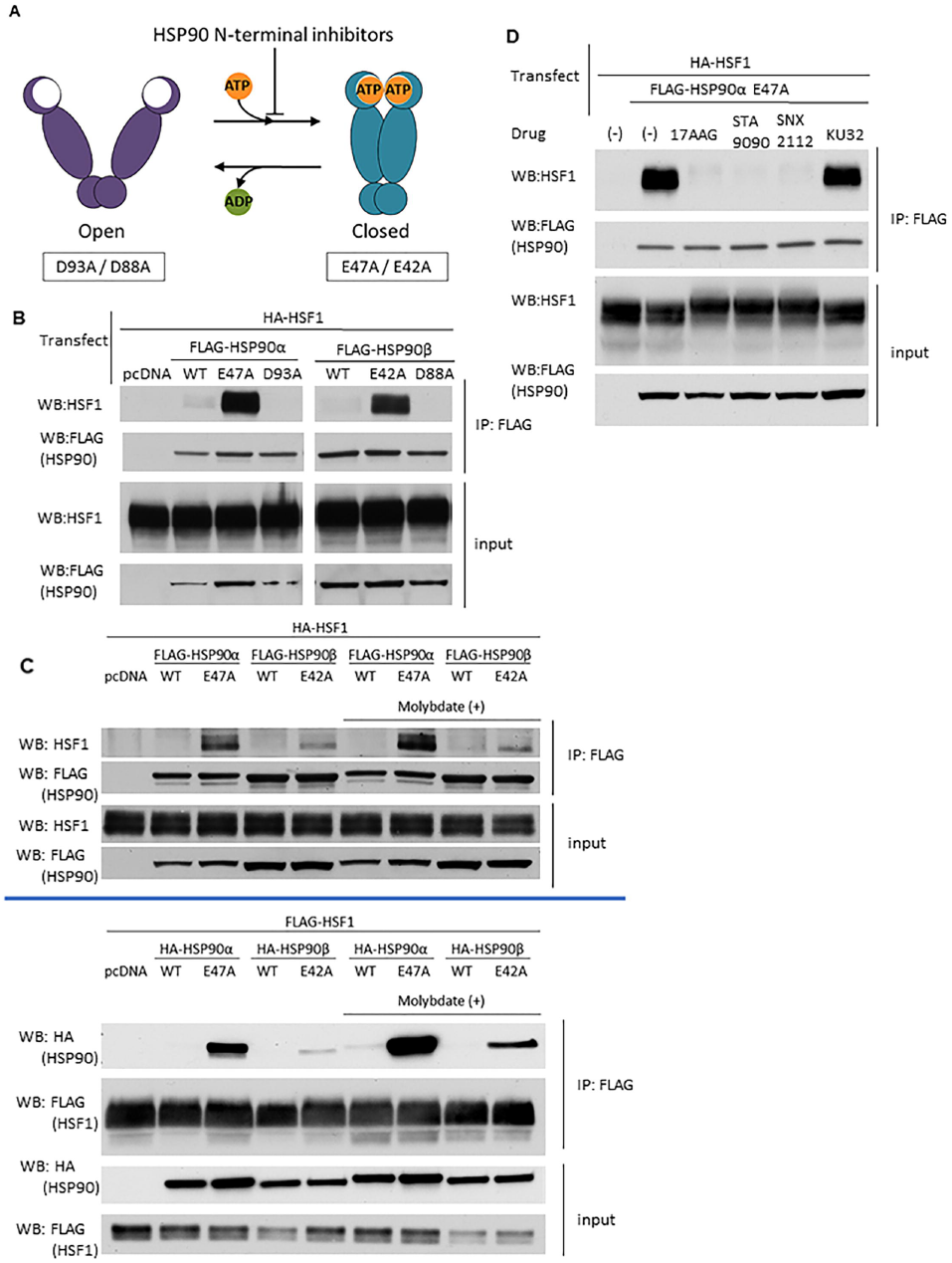
HSF1 is bound by HSP90 in the “closed” conformation. A. HSP90 ATP-driven conformational cycle. HSP90α E47A/HSP90β E42A mutants bind ATP but cannot hydrolyze it and remain trapped in the “closed” N-domain dimerized conformation. Conversely, HSP90α D93A/HSP90β D88A mutants do not bind ATP and remain trapped in an “open” N-domain undimerized conformation. B. HSF1 interaction with HSP90 “closed” and “open” mutants. HEK293 cells were co-transfected with HA-HSF1 and with each FLAG-HSP90 construct, harvested and analyzed for HSF1/HSP90 association by anti-FLAG IP - HSF1 WB. Pull-down experiments were repeated at least twice. C. Reproducible and reciprocal HSF1 interaction with “closed” HSP90α E47A and HSP90β E42A mutants in the presence and absence of MoO4. Interactions were assessed by anti-FLAG (HSP90) IP - HSF1 WB (top panel) or anti-FLAG (HSF1) IP – HA (HSP90) WB (bottom panel). Pull-down experiments were repeated at least twice. D. Effect of HSP90 inhibitors on HSF1-HSP90α E47A interaction. HEK293 cells were transfected with HA-HSF1 along with FLAG-HSP90α E47A, treated with N-terminal or C-terminal HSP90 inhibitors for 4 hours and analyzed by anti-FLAG (HSP90) IP-HSF1 WB. Pull down experiments were repeated at least twice. Full blots used to create cropped figure panels are shown in Supplementary Figure 6.

Mutation of key residues within the N-terminal domain of HSP90 stabilize either the “closed” (HSP90α E47A, HSP90β E42A; allow ATP binding but prevent hydrolysis) or “open” (HSP90α D93A, HSP90β D88A; prevent ATP binding) conformations. We co-transfected HEK293 cells with plasmids encoding FLAG-tagged wild type or conformationally restricted HSP90 proteins together with wild type HA-HSF1, harvested the cells the following day, and observed the protein-protein interactions by anti-FLAG immunoprecipitation-western blot (IP-WB) analysis (Figure 1B, C). The relative expression of HSF1 in transfected and untransfected cells is shown in Supplemental Figure 1A. We found that wild type HSF1 strongly interacts, without the need for crosslinking, with both “closed” HSP90 mutants but not with the pair of “open” HSP90 mutants. Importantly, the results with transfected HSF1 were confirmed by co-immunoprecipitation of “closed” HSP90 with endogenous HSF1 in HEK293 cells (Supplemental Figure 1B). As previously reported ^7^, interaction of wild type HSP90, which samples all available conformational states, with full-length HSF1 is not detectable in the absence of protein crosslinking. This result is consistent with the observation that human HSP90, in contrast to the yeast chaperone, is most likely to be found in the “open” conformation, even when ATP is bound^37^. Moreover, we observed that HSP90α E47A bound more avidly to HSF1 than did HSP90β E42A, suggesting that the stress-inducible HSP90α isoform may be a stronger negative regulator of HSF1 activity compared to the constitutively expressed HSP90β isoform, as previously proposed ^36^.

To further assess the interaction between wild type HSF1 and the “closed” HSP90 mutants, we repeated the reciprocal experiment in the absence or presence of the anion molybdate (MoO_4_) Molybdate has been reported to stabilize a high affinity client binding state of HSP90^38^. Target proteins were pulled down via either FLAG-HSP90 or FLAG-HSF1 (Figure 1C). Although, the HSF1/ “closed” HSP90 interaction was confirmed, molybdate only moderately increased the interaction of the “closed” HSP90 mutants with no discernable effect on interaction of wild type HSP90 with HSF1, indicating that HSP90 interacts with HSF1 in a manner that is likely unique when compared to other clients such as protein kinases ^38 39^.

Administration of HSP90 inhibitors often results in apparent induction or augmentation of the HSR. Currently, there are two general classes of inhibitors. The more clinically advanced N-terminal inhibitors competitively bind to the ATP pocket and prevent or disrupt nucleotide-dependent N-domain dimerization^40^. These inhibitors include 17AAG (tanespimycin), STA9090 (ganetespib) and SNX2112, among others ^30^. A second class of inhibitors, based on the antibiotic novobiocin, binds a putative nucleotide-binding domain in the C-terminal portion of HSP90 (adjacent to the C-terminal dimerization domain) and inhibits the chaperone in a distinct manner^41^. The synthetic novobiocin derivative KU32 is representative of this class of inhibitors^42^. To explore how N- and C-terminal inhibitors might affect HSF1/HSP90 interaction, we co-transfected HEK293 cells with wild type HA-HSF1 and FLAG-HSP90α E47A, briefly treated cells with inhibitors the following day, harvested and analyzed protein-protein interactions by IP-WB (Figure 1D, Supplemental Figure 1C). Each of the three N-terminal inhibitors disrupted the interaction between HSF1 and HSP90α E47A, consistent with their disruption of N-domain dimerization. In contrast, KU32 did not disrupt the interaction between HSF1 and “closed” HSP90, indicating that this is a unique property of ATP-competitive, N-terminal HSP90 inhibitors.

### Mapping of HSP90 and HSP70 interactions across HSF1

To investigate the interactions of HSP90 and HSP70 with HSF1 in the absence of a crosslinking agent, we constructed and transiently expressed in HEK293 cells a series of HSF1 truncation and internal deletion mutants, and we analyzed them for chaperone interaction by IP-WB (Figure 2A and B). Of note, N195, an N-terminal truncation mutant lacking the DBD and most of the HR-A/B domain, expressed very poorly (see blot in Figure 2B) and data obtained using this mutant should be considered in this context. These HSF1 constructs were used to initially query the interaction of HSP90 and HSP70 across the TAD, HR-C, RD, HR-A/B and DBD domains of HSF1 (Supplemental Figure 2A, B).

**Figure 2:**
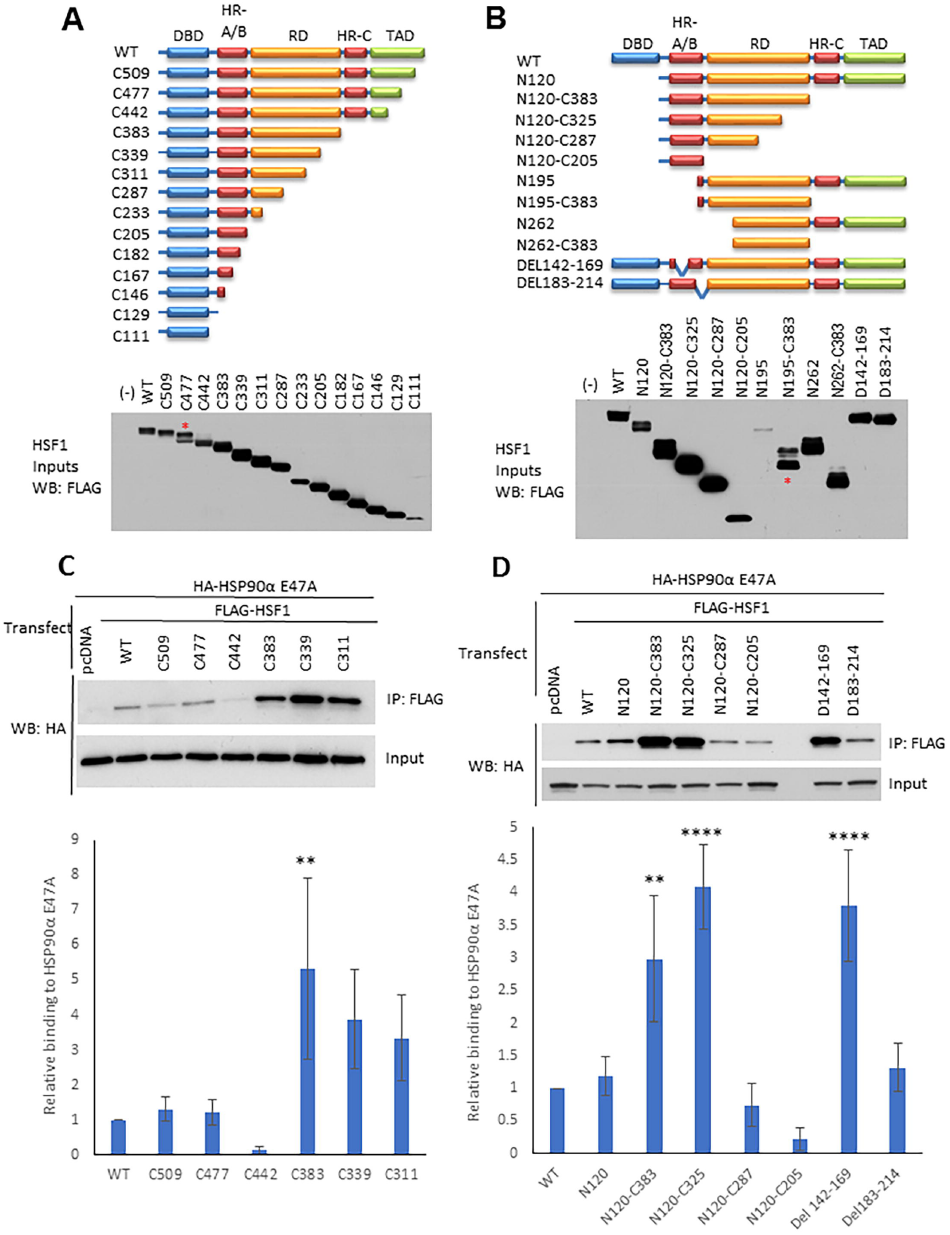
Mapping interaction sites of “closed” HSP90 on HSF1. A. Schema of FLAG-HSF1 C-terminal truncation constructs and their expression in HEK293 cells. The associated blot shows expression of each construct in HEK293 cells. Equal amounts of DNA were used for transfection. For construct C477, the upper band represents the intact truncation (designated by an asterisk), while the lower band is likely a proteolytic fragment. B. Schema of FLAG-HSF1 N-terminal truncation and internal deletion constructs and their expression in HEK293 cells. The associated blot shows expression of each construct in HEK293 cells. Equal amounts of DNA were used for each transfection. Note the poor expression of N195. For N195-C383, the lower band represents the intact construct (designated by an asterisk); the upper band may reflect a post translationally modified peptide. C. Interaction of selected FLAG-HSF1 C-terminal truncation constructs with HA-tagged HSP90α E47A. HEK293 cells were transfected with either FLAG-HSF1 constructs or FLAG-pcDNA control and HA-HSP90α E47A, and then analyzed by anti-FLAG (HSF1) IP followed by HA (HSP90α E47A) WB. Pull downs were performed in triplicate and western blots analyzed by densitometry. All HA IP bands were normalized to their respective FLAG IP signals (see Supplemental Figure 6, compilation of full blots) and then normalized to the WT signal. A one-way ANOVA utilizing multiple comparisons was performed to determine significant differences in HSP90 interaction between truncated HSF1 constructs and F-HSF1 WT. A representative blot is shown above the bar graph. Significance is indicated by asterisks (** indicates p< 0.01). The p-value for the difference between C339 and WT interaction was 0.0534. D. Interaction of FLAG-HSF1 N-terminal truncation and internal deletion constructs with HA-tagged HSP90α E47A: HEK293 cells were transfected and analyzed as in panel C. Significance is indicated by asterisks (** indicates p< 0.01 and **** indicates p< 0.0001). Full blots used to create cropped figure panels and for densitometry analysis for the bar graphs are shown in Supplementary Figure 6.

Based on these preliminary data obtained with the closed HSP90 mutants, a subpanel of HSF1 truncation and deletion mutants were selected to assess association of HSP90α E47A. Pulldowns were repeated in triplicate, resulting blots were subjected to densitometry to generate mean values +/- standard deviations and p values to assess significance (compared to association of HSP90α E47A with full length HSF1) were obtained. These data are shown in Figure 2, panels C and D. Analysis of HSF1 C-terminal truncations revealed a distinct interaction motif for “closed” HSP90 E47A (Figure 2C). The “closed” mutant bound most intensely to HSF1 proteins containing the HR-A/B domain and a portion of the heat-sensing RD, but lacking the HR-C and TAD. The clear difference in HSP90 interaction between the C442 and C383 HSF1 truncation mutants suggests that presence of the HSF1 HR-C + TAD domain inhibits HSP90 interaction. Although HSP90 E47A bound weakly to more extensive C-terminal truncations, binding was not significantly different from wild type HSF1. The dual importance of both the HR-A/B and RD domains for HSP90 binding is supported by the observation that neither the isolated HR-A/B domain (N120-C205, Figure 2D and Supplemental Figure 2B) nor the isolated RD domain (N262-C383, Supplemental Figure 2B) display binding to HSP90.

Importantly, deletion of the HR-C domain dramatically improves the binding of both “closed” and endogenous HSP90, suggesting that HR-C intermolecular binding to HR-A/B (characteristic of inactive HSF1 monomers) interferes with HSP90 binding and providing further evidence that is inconsistent with a role for HSP90 in sequestering monomeric HSF1. Finally, the HSF1 HR-A/B internal deletion construct DEL142-169, where amino acids 142-169 were removed, bound “closed” HSP90 while the HR-A/B internal deletion construct DEL183-214 did not (Figure 2D and Supplemental Figure 2D), suggesting that the C-terminal half of the HR-A/B is necessary for HSP90 binding.

While the binding pattern of endogenous HSP90 generally mirrors that of “closed” HSP90 (although, in contrast to “closed” HSP90, detectable binding of endogenous HSP90 requires deletion of the HR-C domain, see Supplemental Figure 2A, B), endogenous HSP90 also uniquely binds the HSF1 constructs N195 and N262 but not the equivalent HSF1 mutants lacking the HR-C and TAD domains (N195-C383 or N262-C383, see Supplemental Figure 2B). These findings are consistent with the interaction of endogenous HSP90 (primarily existing in an N-domain undimerized conformation) with the HR-C domain, as recently reported by Mayer and colleagues for HSP90β. Importantly, these authors demonstrated that HSP90 binding to the HSF1 HR-C domain promotes its temperature-sensitive dissociation from HR-A/B, thus facilitating HSF1 trimerization^25^. Like our data, these findings are also not consistent with a role for HSP90 in suppressing HSF1 trimerization.

In contrast to the binding profile of HSP90, endogenous HSP70 interacts robustly with full-length HSF1 as well as with most of the C-terminal truncation and deletion mutants (Supplemental Figure 2A and B), with the exception of those mutants containing increasingly truncated RD. Endogenous HSP70 also bound poorly to N120-C205 (isolated HR-A/B domain) despite ample expression of this HSF1 protein fragment. Taken together, these data identify the RD as being sufficient for HSP70 binding in unstressed cells, although HSP70 binding to other domains of HSF1 has also been reported ^26 27^.

To further compare the role of the HR-A/B domain in HSF1 trimerization and in HSP90 interaction, we made a set of alanine scanning mutants (145A_4_, 151A_5_, 161A_5_) between amino acids142-169 (see Methods) and compared them in the experiments below with the internal deletion constructs DEL142-169 and DEL183-214. Using IP-WB analysis of co-transfected WT HA-HSF1 and mutant FLAG-HSF1, we observed that the HR-A/B domain alanine mutants did not disrupt HSF1 intermolecular interaction, in distinct contrast to the HR-A/B deletion mutants DEL142-169 and DEL182-214 (Supplemental Figure 2C). In contrast, these alanine mutants bound “closed” HSP90α E47A to a greater extent compared to WT HSF1 (but similarly to DEL142-169 (Supplemental Figure 2D and Figure 2D). These data are consistent with the hypothesis that the HR-A/B region contributing to HSF1 trimerization is distinct from the HR-A/B region that is required for HSP90 binding. Further, our findings emphasize that the presence or absence of HSP90 binding has no impact on HSF1 trimerization (see Figure 3B, D and Supplemental Figure 5). Finally, the enhanced HSP90 binding of the HSF1 truncation and deletion mutants shown in Figure 2C, D does not involve possible heterotrimerization of the transfected HSF1 constructs with endogenous HSF1, as similar results are obtained when the HSF1 truncation mutants are transfected into HEK293 cells stably knocked down for endogenous HSF1 (Supplemental Figure 2E).

**Figure 3:**
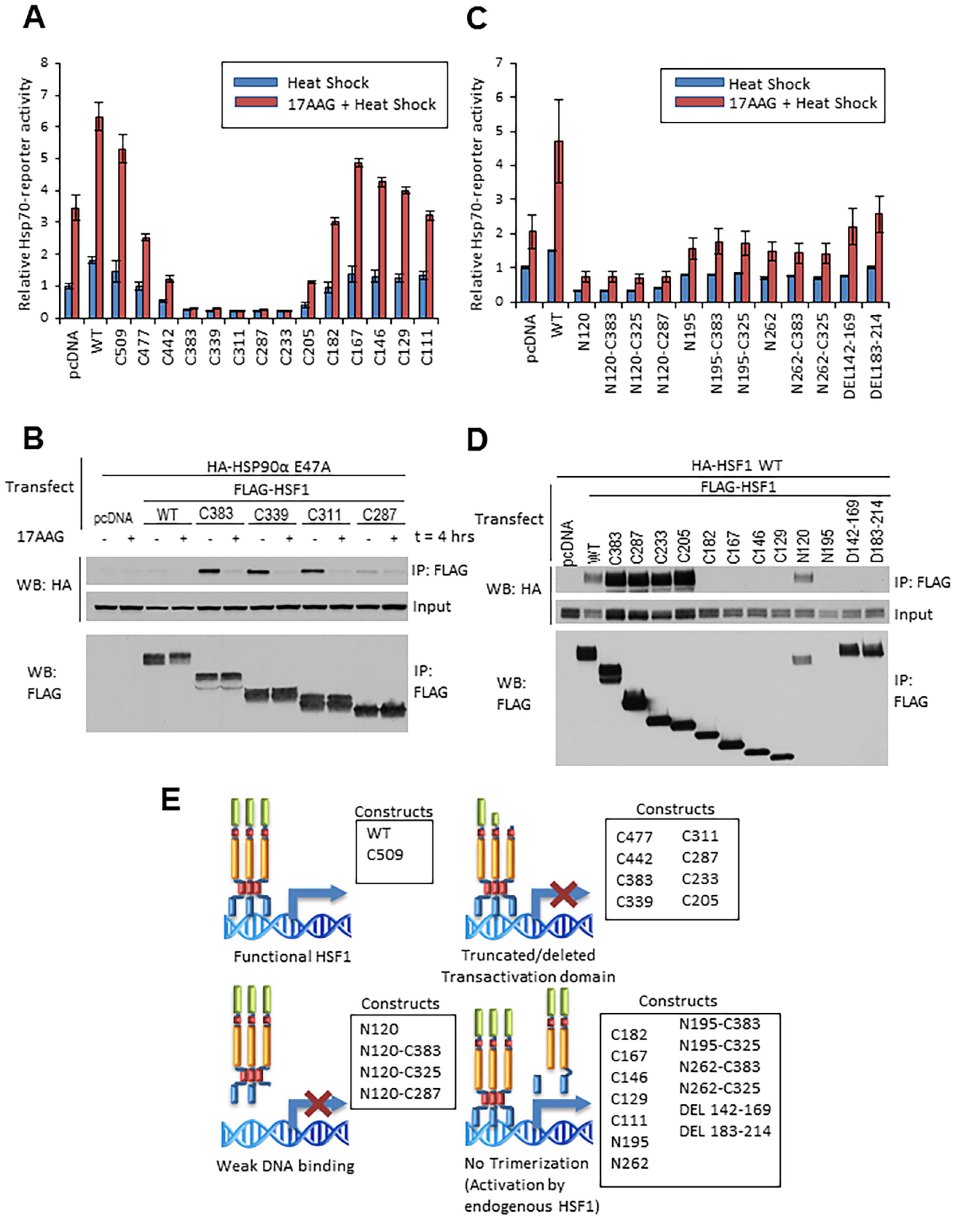
Effects of HSF1 truncation and internal deletion mutants on hsp70 promoter reporter activity and HSF1 dimerization. A.Expression of TAD-deleted HSF1 constructs inhibits endogenous HSF1 driven *hsp70* promoter-reporter activity, while removal of the HR-A/B domain abrogates dominant negative activity. HEK293 cells were transfected with *hsp70* promoter-reporter construct and HSF1 C-terminal truncation constructs, treated with DMSO or 17AAG for 1 hour, heat shocked at 42°C for 30 minutes and allowed to recover for 4 hours before harvesting. B. Disruption of HA-tagged HSP90α E47A interaction with HSF1 C-terminal RD truncations by 17AAG. HEK293 cells were transfected with HA-HSP90α E47A and FLAG-HSF1 RD truncation mutants, then analyzed by anti-FLAG (HSF1) IP followed by HA (HSP90α E47A) WB. Pull-down experiments were repeated at least twice. C. Deletion of the DBD and retention of the HR-A/B domain inhibits *hsp70*-reporter activity driven by endogenous HSF1. HEK293 cells were transfected, treated and processed as described in panel A. D. Deletion of the HR-A/B domain disrupts HSF1 dimerization. HEK293 cells were transfected with WT HA-HSF1 and FLAG-HSF1 deletion constructs, then analyzed by anti-FLAG IP and HA WB. Pull-down experiments were repeated at least twice. E.Summary of the effects of HSF1 domain deletions on *hsp70*-reporter activity. Full blots used to create cropped figure panels are shown in Supplementary Figure 6.

### The TAD and DBD are essential for HSF1 transcriptional activity while the HR-A/B domain is required for dominant negative activity of truncated HSF1

To further explore potential relationships between the ability of HSF1 to (1) form intermolecular trimers, (2) activate transcription, and (3) interact with HSP90, we examined the impact of HSF1 N/C-truncations and internal deletions on *hsp70* promoter-reporter activity as a readout of the HSR, and we complemented these observations with IP-WB analysis of HSF1 trimerization and HSF1-HSP90 interaction. Overexpression of WT HSF1 increased *hsp70*-reporter activity in response to heat shock by nearly 2-fold above a pcDNA control. Brief pretreatment of cells with 17AAG prior to heat shock increased *hsp70*-reporter activity in pcDNA control by 3.5-fold compared to heat shock alone, and a similar fold increase in reporter activity was observed upon 17AAG exposure prior to heat shock in cells transfected with WT HSF1 (Figure 3A). For reference, reporter activity in response to 17AAG alone, compared to heat shock and heat shock+ 17AAG is shown in Supplemental Figure 3A. Both heat-induced and 17AAG + heat-induced *hsp70*-reporter activity were progressively impacted by increasing truncation of the TAD (C-terminal deletion constructs C509, C477 and C442), consistent with the importance of this domain in modulating HSF1 activation ^43^. Complete removal of TAD, HR-C and RD domains (constructs C383 through C205) fully repressed *hsp70*-reporter activity even with 17AAG pretreatment, suggesting that the presence of the HR-A/B domain of these mutants is essential for dominant-negative activity, as a consequence of disrupting endogenous HSF1 trimers. Indeed, truncation through the C-terminal portion of the HR-A/B domain (C182) restored *hsp70*-reporter response to that seen after treating pcDNA control cells with heat and heat + 17AAG. Consistent with this result, HSF1 C182 is unable to form an intermolecular interaction with wild type HSF1, as are additional HR-A/B truncations (Figure 3D). Unexpectedly, further truncation (C167, C146 and C129) through the HR-A/B, increased *hsp70*-reporter activity above that of the pcDNA control, suggesting the presence of a repressing motif located between amino acids 167 and 182 of HSF1 (Figure 3A).

Our previous observation of HSP90 binding to a series of RD-containing C-terminal truncation mutants (C383, C339, C311, Figure 2C) suggested that HSP90 binding might be involved in repressing HSF1-driven *hsp70*-reporter activity. To address this possibility, we used IP-WB to investigate the effect of 17AAG on the interaction of HSP90α E47A with these HSF1 RD truncation mutants since these mutants repressed *hsp70*-reporter activity even in the presence of 17AAG (Figure 3A). Consistent with the data in Figure 2C, we observed increased binding of HSP90α E47A to the RD truncation mutants C383, C339 and C311 when compared to wild type HSF1. However, upon addition of 17AAG, HSF1-HSP90α E47A interaction was fully disrupted (Figure 3B), while *hsp70*-reporter activity remained repressed (Figure 3A). These data indicate that robust HSP90 interaction with HSF1 is not responsible for the dominant negative activity of these mutants. Consistent with this hypothesis, the RD truncation mutant C287 bound poorly to HSP90αE47A, yet displayed dominant negative activity (see Figures 3A and B). Taken together, our findings support the hypothesis that the dominant negative activity of HSF1 C-terminal truncations retaining an intact HR-A/B domain is due to disruption of endogenous full length HSF1 trimers necessary for transcriptional activity.

Next, we assessed the effects of another set of HSF1 N/C-terminal deletion constructs on *hsp70*-reporter activity (Figure 3C). Here we observed that the deletion constructs N120, N120-C383, N120-C325 and N120-C287, all lacking the DBD but including an intact HR-A/B domain, repressed *hsp70*-reporter activity below the level seen with empty plasmid, consistent with dominant negative inhibition of endogenous HSF1. Removal of the HR-A/B domain (N195, N262) relieved this repression to some extent, even when the TAD and RD were removed (N195-C383, N195-C325, N262-C383, N262-C325). Finally, the internal HR-A/B deletion mutants DEL142-169 and DEL182-214 supported *hsp70*-reporter activity at a level similar to that of empty plasmid, confirming that an intact HR-A/B domain is required for dominant negative activity. The statistical significance of the changes in reporter activity can be found in Supplemental Figure 3B. Of note, these data are in contrast to the impact of these two HR-A/B internal deletions on HSP90 binding (Figure 2D, right two lanes).

To profile the ability of these deletion constructs to affect HSF1 intermolecular interaction (indicative of trimerization capability), we analyzed by IP-WB the interaction of co-transfected wild type HA-HSF1 with various FLAG-HSF1 deletion mutants. We found that FLAG-tagged deletion mutants (C383, C287, C233, C205 and N120) that dominant negatively repressed *hsp70*-reporter activity interacted efficiently (equivalent or enhanced interaction compared to FLAG-tagged wild type HSF1) with wild type HA-HSF1. In contrast, deletion mutants (C182, C167, C146, C129, DEL142-169, DEL183-214) that did not strongly repress *hsp70*-reporter activity did not interact with WT HA-HSF1 (Figure 3D). We excluded N195 from this analysis due to its poor expression as shown in Figures 2B and 3D. Likewise, the dominant negative HSF1 mutants N120, N120-C383, N120-C325 and N120-C287 strongly interacted with full length HSF1, while N195-C383 (which showed no dominant negative activity) did not (Supplemental Figure 3C and D). Combined with our previous observations, these data further confirm that HSF1 mutant-mediated dominant negative repression of endogenous HSF1-driven transcription depends on the ability to form an intermolecular interaction with wild type HSF1. A cartoon summarizing these data is provided in Figure 3E.

### Impact of RD phosphorylation state on HSP90 inhibitor effect on HSF1 activity and mobility shift in SDS-PAGE, and on HSF1 interaction with HSP90

Stress-inducible phosphorylation of the HSF1 RD coincides with acquisition of transactivation capacity, although a functional requirement of RD phosphorylation for HSF1 activation has not been demonstrated. Indeed, a recent report dissociates RD phosphorylation from HSF1 activation and suggests that RD phosphorylation may serve as a rheostat to modulate the degree of activation^23^. Since N-terminally targeted HSP90 inhibitors also cause a distinct,phosphorylation-dependent mobility shift of HSF1 coincident with disrupting its association with conformationally restricted HSP90 proteins (see Figure 1D, Supplemental Figure 1C and Supplemental Figure 4), we wished to determine whether these drugs promoted phosphorylation of sites only within the RD and whether RD phosphorylation was a prerequisite for HSP90 inhibitor-enhanced HSF1 activity. For this purpose, we used the delta PRD (dPRD) HSF1 construct previously described, in which all phosphorylation sites within the RD have been mutated to alanine (see Figure 4A)^23^. Using the *hsp70*-reporter assay, we confirmed that dPRD HSF1 was more sensitive to heat shock, and to the combination of heat shock and 17AAG, compared to wild type HSF1 (Figure 4B). Unexpectedly, the mobility of HSF1 dPRD was also shifted when challenged with three N-domain HSP90 inhibitors (Figure 4C), strongly suggesting that HSP90 inhibitor-induced HSF1 phosphorylation is not restricted to site(s) within the RD (Supplemental Figure 4). Importantly, lack of phosphorylation within the RD domain does not affect trimerization of either full length HSF1 or the various HSF1 deletion mutants discussed above (Supplemental Figure 3E).

**Figure 4:**
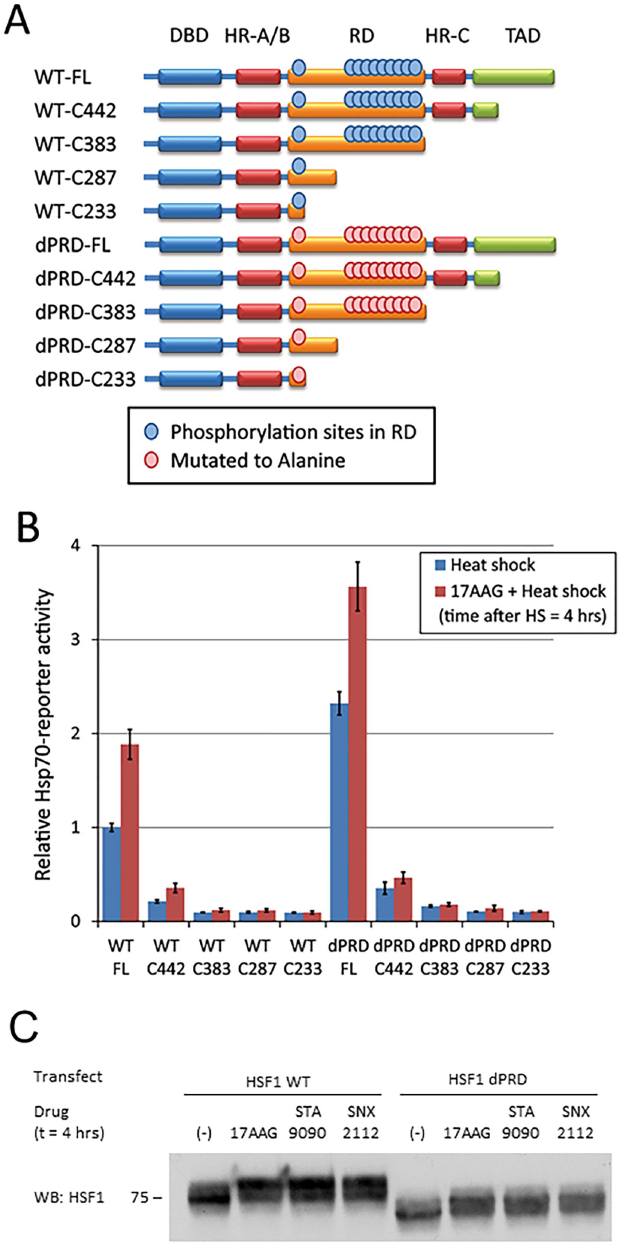
Impact of RD phosphorylation on HSP90 inhibitor effect on HSF1 activity and mobility shift in SDS-PAGE, and on HSF1 interaction with HSP90. A. Schema of WT and dPRD FLAG-HSF1 constructs depicting RD phosphorylation sites mutated to alanine in dPRD-HSF1. B. The N-terminal HSP90 inhibitor 17AAG enhances transcriptional activity of dPRD HSF1. HEK293 cells were transfected with *hsp70*-reporter construct and HSF1 C-terminal truncation constructs, treated with DMSO or 17AAG for 1 hour, heat shocked at 42°C for 30 minutes and allowed to recover for 4 hours before harvesting. This experiment was repeated three times and values represent mean +/- S.D. C. N-terminal HSP90 inhibitors induce a mobility shift in dPRD HSF1 similarly to that seen with WT-HSF1. HEK293 cells were transfected with WT or dPRD HSF1, treated with the three N-terminal HSP90 inhibitors as shown, and harvested and HSF1 was analyzed by WB. Full blots used to create cropped figure panels are shown in Supplementary Figure 6.

### HSP90 inhibition increases duration of HSF1 binding to *hsp70* promoter and prolongs duration of HSF1 transcriptional activity in response to heat shock

To assess the outcome of combining HSP90 inhibition with heat shock compared to either inhibitor or heat shock alone, we monitored the extent and duration of HSF1 occupancy of the *hsp70* promoter by chromatin immunoprecipitation (ChIP) with antibody to HSF1. HEK293 cells were pretreated for 1 hour at 37°C with 10 μM 17AAG or left untreated, and then exposed to heat shock (42°C for 30 minutes). After recovery for varying times at 37°C (no recovery, 2 hours,4 hours, 6 hours), HSF1 ChIP was performed and pulldown of associated *hsp70* promoter sequences was assessed by PCR. An additional group exposed to 17AAG but in the absence of heat shock was also subjected to ChIP analysis at the indicated times (based on recovery time after heat shock). The top panel in Figure 5A shows a representative experiment in which the expected PCR fragments were run on an agarose gel. In the bottom panel, PCR data reflecting the extent and duration of HSF1 occupancy of the *hsp70* promoter were quantified by densitometry (3 independent experiments +/- standard deviation), normalized to input and graphed as shown (Figure 5A, values obtained in the absence of both heat shock and 17AAG pretreatment were arbitrarily set to 1). In the absence of 17AAG, HSF1 binding to the *hsp70* promoter was highest immediately after heat shock (no recovery). Promoter occupancy declined below baseline by 2 hours and remained below baseline for the duration of the experiment. In contrast, 1-hour pre-exposure of cells to 17AAG prior to heat shock resulted in an increased duration of HSF1 binding to the *hsp70* promoter (significantly above baseline at 2 hours of recovery and significantly above the heat shock only value at later time points). Importantly, the 17AAG alone samples displayed the same maximal level of HSF1 binding to *hsp70* promoter as did both the heat shock alone samples and the heat shock + 17AAG samples. However, in contrast to cells treated with heat shock alone, HSF1 promoter occupancy did not decline as rapidly in cells treated with 17AAG alone or in combination with heat shock. Thus, the level of HSF1 binding to DNA in these groups remained significantly above heat shock only values for the full 6-hour recovery period (at least 4 hours longer than after exposure to heat shock alone).

Additionally, we assessed HSF1 transcriptional activity in HEK293 cells after heat shock alone, 17AAG alone, or after pre-exposure to 17AAG followed by heat shock, using the same experimental conditions as for the ChIP analysis above (Figure 5B). For this purpose, we examined endogenous *HSP70* gene expression by RT-qPCR. Similar to the impact of heat shock on HSF1 promoter occupancy, relative *HSP70* mRNA expression peaked after 2 hours of recovery from heat shock and declined thereafter to below baseline by 6 hours. In contrast, 17AAG alone promoted a modest increase in *HSP70* mRNA (although below the peak seen after heat shock) that remained elevated at 2 and 6 hours. After combined 17AAG treatment and heat shock, *HSP70* mRNA level increased on par with exposure to heat shock alone (similar values at 2 hours), but continued to increase to a peak value at 4 hours (25 % higher than the heat shock alone peak value) that was maintained at 6 hours. Importantly, the 4- and 6-hour increase in *HSP70* mRNA after combination treatment was significantly higher (2 – 2.5-fold) compared to HSP90 inhibitor alone.

**Figure 5:**
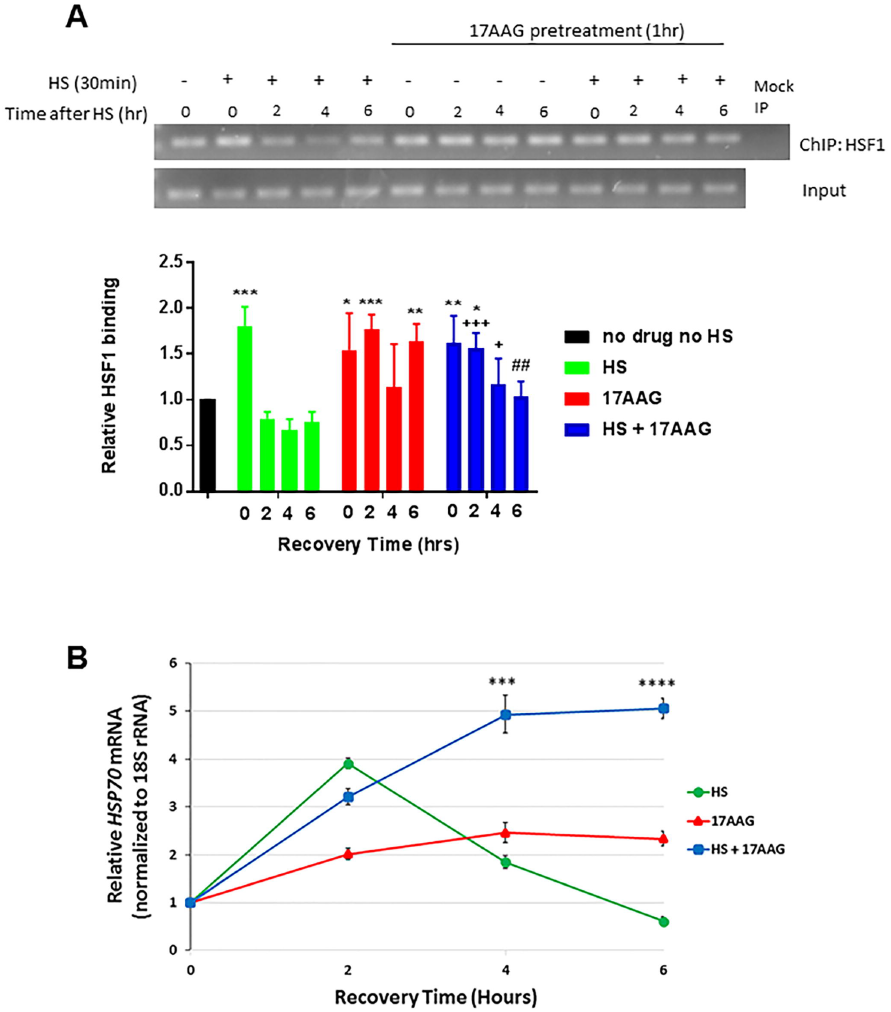
HSP90 inhibition increases duration, but not extent, of HSF1 binding to the hsp70 promoter and enhances HSF1 transcriptional activity in response to heat shock. A.Chromatin immunoprecipitation using HSF1 antibody followed by PCR amplification of the *hsp70* promoter sequence (top, representative DNA gel; bottom, graphical display of band optical density data from 3 independent experiments +/- S.D.). HEK293 cells were either subjected to heat shock (42°C/30 minutes), treated with 17AAG, or both. The 17AAG groups were pre-exposed to the HSP90 inhibitor (10 μM, 1 hour) prior to heat shock. Samples were then put back in a 37°C incubator to recover for the indicated times. Values for the’ no drug, no HS’ control A.(first lane) were set to 1, and remaining values are expressed as fold above or below this baseline value. This experiment was repeated three times and values represent mean +/- S.D. A two-way ANOVA was used to analyze the samples for statistically significant differences compared to other samples at a given time. Significant difference compared to’ no drug, no HS’ is indicated by asterisks (*** indicates p< 0.001, ** indicates p< 0.01, and * indicates p< 0.05);’ 17AAG + HS’ is also significantly lower than’ 17AAG’ alone at 6hr (## - p< 0.01) and significantly higher than’ HS’ alone at 2 and 4hr (+++ indicates p< 0.001, + indicates p< 0.05). B.Endogenous HSP70 mRNA was quantified by RT-qPCR in HEK293 cells subjected to heat shock alone, 17AAG alone, or heat shock after pre-exposure to 17AAG, using identical experimental conditions as in panel A. This experiment was performed in triplicate and values represent mean +/- S.D. A two-way ANOVA was used to analyze for statistically significant differences. Notably, there was a statistically significant difference between’ 17AAG alone’ and’ 17AAG + HS’ samples at 4 and 6 hours (*** indicates p< 0.001 and **** indicates p< 0.0001).

## Discussion

In this study, we assessed the importance of HSP90 conformation for binding to HSF1 and we found that only the ATP-dependent “closed” (e.g., N-domain dimerized) conformational mutants E47A (HSP90α) and E42A (HSP90β) readily co-precipitate full-length HSF1, with HSP90α displaying the stronger interaction. As HSP90α is stress-inducible and transcriptionally regulated by HSF1, this provides an appealing feedback control mechanism to attenuate the HSR, consistent with a model first proposed nearly twenty years ago^8 44^. Indeed, our previous finding that transient over-expression of HSP90α reduced HSF1 transcriptional activity supports this possibility^36^. In the absence of these conformationally restrictive mutations, mammalian HSP90 only rarely and transiently occupies the “closed” conformation, even in the presence of ATP, with the bulk of the protein at steady-state occupying an N-domain undimerized conformation ^37^. This is inconsistent with a role for HSP90 in binding to and sequestering HSF1 monomers to prevent trimerization, as has also been proposed ^7^.

To identify chaperone binding sites in HSF1, we used a series of Flag-tagged HSF1 truncation and internal deletion mutants to interrogate binding of HSP90 and HSP70, including the binding of transiently expressed HSP90αE47A or HSP90β E42A. Using this approach, we identified a region in HSF1 that associates most strongly with “closed” HSP90 (Supplemental Figure 5). This motif is comprised of the HR-A/B trimerization domain in the context of a portion of the RD. Binding of “closed” HSP90 to this motif is dramatically enhanced upon deletion of the HSF1 HR-C and TAD domains. Since the HR-C domain interacts with the HR-A/B domain to retard HSF1 trimerization under basal conditions, these data are consistent with the preferential binding of “closed” HSP90 to HSF1 trimers. Residues in the C-terminal half of the HR-A/B domain (amino acids 183-214) are necessary and sufficient for binding. Of note, while endogenous HSP90 shares the HR-A/B + RD binding motif, visualization of endogenous HSP90 binding requires deletion of HR-C and TAD domains, very likely due to the weaker affinity of endogenous HSP90 (primarily existing in an “open” conformation) for client interactors. Taken together, these findings are not consistent with a role for HSP90 in repressing HSF1 trimerization.

In contrast to HSP90, endogenous HSP70 binds to the isolated RD as well as other domains in HSF1. Only the isolated HR-A/B and DBD domains display weak interaction with HSP70. These data build upon earlier work by Shi et al demonstrating repression of HSF1 transcriptional activity as a consequence of HSP70 binding to the TAD^26,27^. Our observation that HSP70 also binds to the stress sensing RD may be relevant to the recent proposal that HSP70 provides an on/off switch for dynamic regulation of stress-induced HSF1 activation in yeast ^28^. Whether HSP70 plays a similar role in regulating HSF1 activation in mammalian cells remains to be explored.

We used these HSF1 truncation and deletion mutants to identify the HR-A/B domain as necessary for dominant negative activity of HSF1 mutants lacking the TAD or the DBD. Importantly, and consistent with our model, we found that HSP90 binding to the HR-A/B is not required for dominant-negative activity as it is not reversed by 17AAG. Instead, dominant-negative activity depends on the ability to form an intermolecular interaction with other HSF1 monomers, suggesting that each monomer of an HSF1 trimer must be intact in order to attain optimal HSF1 transcriptional activity.

A panel of clinically evaluated N-terminal HSP90 inhibitors uniformly enhance HSF1 phosphorylation coincident with dissociation of HSF1 from HSP90. One or more drug-sensitive phosphorylation sites must lie outside of the RD, as an HSF1 protein with all putative RD phosphorylation sites mutated to alanine (dPRD HSF1) still underwent a phosphorylation-dependent mobility shift upon HSP90 inhibitor treatment. We are currently exploring the relevance of this observation to the activity of both wild type and dPRD HSF1, as well as the mechanistic impact of HSP90 inhibitor-mediated HSF1 phosphorylation on its transcriptional activity.

HSP90 ATP-competitive N-terminal inhibitors efficiently dissociate HSF1 from HSP90 “closed” mutants, likely as a consequence of ATP displacement and subsequent disruption of N-domain dimerization^40^. If ATP-bound, transiently N-domain dimerized HSP90 (including stress-inducible HSP90α) provides feedback control of the HSR by removing HSF1 trimers bound to heat shock promoter elements, HSP90 inhibition and subsequent dissociation from HSF1 would interfere with this process and would be predicted to prolong and/or intensify the HSR. Consistent with this model, HSF1-CHiP data show that HSP90 inhibition alone and in combination with a brief heat shock significantly enhanced the duration of HSF1 binding to *HSP70* promoter, compared to heat shock alone. When we assessed HSF1 transcriptional activity after these treatments (monitoring endogenous *HSP70* mRNA levels by RT-qPCR), we found that pre-exposure to HSP90 inhibitor markedly enhanced the degree and significantly extended the duration of *HSP70* mRNA increase during recovery from a brief heat shock. HSP90 inhibition alone caused a moderate increase in *HSP70* transcription, likely due to disruption of the dynamic regulation of constitutive HSF1 activity in transformed cells, a phenomenon recently described by Lindquist and colleagues^11 12^.

In summary, this report increases our understanding of the molecular mechanisms that regulate HSF1 and the unique contribution of molecular chaperones, including HSP90 and HSP70, to this process. While not precluding other possibilities, our findings are consistent with a role for HSP90 in attenuating the HSR in an ATP-dependent and conformation-specific manner, a process that is antagonized by ATP-competitive HSP90 inhibitors. A role for HSP90 in disassembling transcriptional regulatory complexes has been described previously ^45^. Our data further suggest that, in contrast to HSP90, the readily observable interaction of endogenous HSP70 with HSF1, and our observation that the RD represents a major interaction motif for this chaperone, support the possibility that HSP70 may provide constitutive, stress-sensitive repression of the HSR in mammalian cells.

## Materials and Methods

### Cell culture

HEK293 cells (American Type Culture Collection) were maintained in DMEM supplemented with 10% FBS. Cells were grown in 6-well or 96-well plates to 50% confluency and then transfected for 16-24 hours with X-tremeGENE 9 transfection reagent (Roche) according to the provided protocol. HEK293 cells stably expressing an shRNA targeting HSF1 was obtained from Dr. Chengkai Dai (Center for Cancer Research, NCI) and has been described previously ^46^.

### Reagents

17AAG was obtained from the Developmental Therapeutics Program, NCI. STA9090 (ganetespib) and SNX2112 were obtained from Synta Pharmaceuticals and Esanex, respectively. The C-terminal inhibitor KU32 was kindly provided by Dr. Brian Blagg (The University of Kansas, Lawrence, KS). MG132 and molybdate were purchased from Sigma-Aldrich. All drugs were used at 10 μM for IP-WB or *hsp70*-reporter assays.

### Plasmids

Sets of ATPase mutants of pcDNA-FLAG-HSP90α/β and pcDNA-HA-HSP90α/β were made by the Quickchange method to alter designated amino acids. The series of C-terminal deletion constructs of pcDNA-FLAG-HSF1 were made by inserting a terminal stop codon into designated regions by the Quickchange method. The series of N-terminal deletion constructs of pcDNA-FLAG-HSF1 were made by ligating truncated HSF1 insert with pcDNA-FLAG backbone by Gibson assembly according to the provided protocol (New England Biolabs).

### Immunoprecipitation and western blotting

Cells were lysed with TGNET buffer [50 mM Tris HCl pH7.5, 5% Glycerol, 100 mM NaCl, 2 mM EDTA, 0.5% Triton X-100] containing protease inhibitor cocktail and phosphatase inhibitor cocktail (Roche). Lysates were centrifuged at maximum speed for 15 minutes at 4°C. Supernatants were subjected to BCA protein assay for normalization, and then 40 μg of protein were used as input lysate to confirm protein expression. From remaining lysates, 700 μg of protein were subjected to immunoprecipitation by addition of 40 μl of Anti-FLAG M2 affinity beads (Sigma-Aldrich, A2220) or HA affinity beads (clone HA-7, Sigma-Aldrich, A2095) followed by incubation with rotation for 2 hours at 4°C. After centrifugation, beads were washed 4 times with TGNET buffer. The proteins were then eluted with 30 μl of 2X SDS sample buffer by boiling at 95°C for 5 minutes. Subsequently, samples were subjected to SDS-PAGE followed by western blotting. The following antibodies were used for western blot: HSF1 (Santa Cruz, sc-13516), HSP90 (Enzo Life Science, ADI-SPA-835), HSP70 (Enzo Life Sciences, ADI-SPA-810), HA (Rockland, 600-401-384), and FLAG (Sigma-Aldrich, A8592).

Additional Methods may be found in Supplemental Information. Materials and data generated in this study are available upon request.

## Acknowledgments

This study was supported by funds from the Intramural Research Program, National Cancer Institute, Center for Cancer Research (L.N.). T.K was supported by a JSPS Research Fellowship for Japanese Biomedical and Behavioral Researchers at NIH. We dedicate this paper to the memory of Dr. Susan Lindquist, a friend and a pioneer in the study of molecular chaperones.

## Author Contributions

T.K. designed experiments, performed experiments and helped write the paper; T.L.P. designed experiments, performed experiments, interpreted the data and helped write the paper; M.L.T., K.H.Y., H.S., K.B. performed experiments and interpreted the data; S.L. and J.B.T. performed experiments, interpreted the data and helped write the paper; M.A.B. and L.S. provided reagents, interpreted data and helped write the paper; H.W. and S.C. interpreted data and helped write the paper; L.N. oversaw the whole project, devised experiments, interpreted data and wrote the paper.

## Data Availability

Materials and data generated in this study are available upon request to the corresponding authors (T.L.P. and L.N.).

